# GIV/Girdin, a Non-receptor Modulator for Gαi/s, Regulates Spatiotemporal Signaling during Sperm Capacitation and is Required for Male Fertility

**DOI:** 10.1101/2021.05.06.442927

**Authors:** Sequoyah Reynoso, Vanessa Castillo, Gajanan D. Katkar, Inmaculada Lopez-Sanchez, Sahar Taheri, Celia R. Espinoza, Christina Rohena, Debashis Sahoo, Pascal Gagneux, Pradipta Ghosh

## Abstract

For a sperm to successfully fertilize an egg, it must first undergo capacitation in the female reproductive tract, and later undergo acrosomal reaction (AR) upon encountering an egg surrounded by its vestment. How premature AR is avoided despite rapid surges in signaling cascades during capacitation remains unknown. Using a combination of KO mice and cell-penetrating peptides, we show that GIV (CCDC88A), a guanine nucleotide-exchange modulator (GEM) for trimeric GTPases, is highly expressed in spermatocytes and is required for male fertility. GIV is rapidly phosphoregulated on key tyrosine and serine residues in human and murine spermatozoa. These phosphomodifications enable GIV-GEM to orchestrate two distinct compartmentalized signaling programs in the sperm tail and head; in the tail, GIV enhances PI3K→ Akt signals, sperm motility and survival, whereas in the head it inhibits cAMP surge and premature AR. Furthermore, GIV transcripts are downregulated in the testis and semen of infertile men. These findings exemplify the spatiotemporally segregated signaling programs that support sperm capacitation and shed light on a hitherto unforeseen cause of infertility in men.

**GRAPHIC ABSTRACT:** 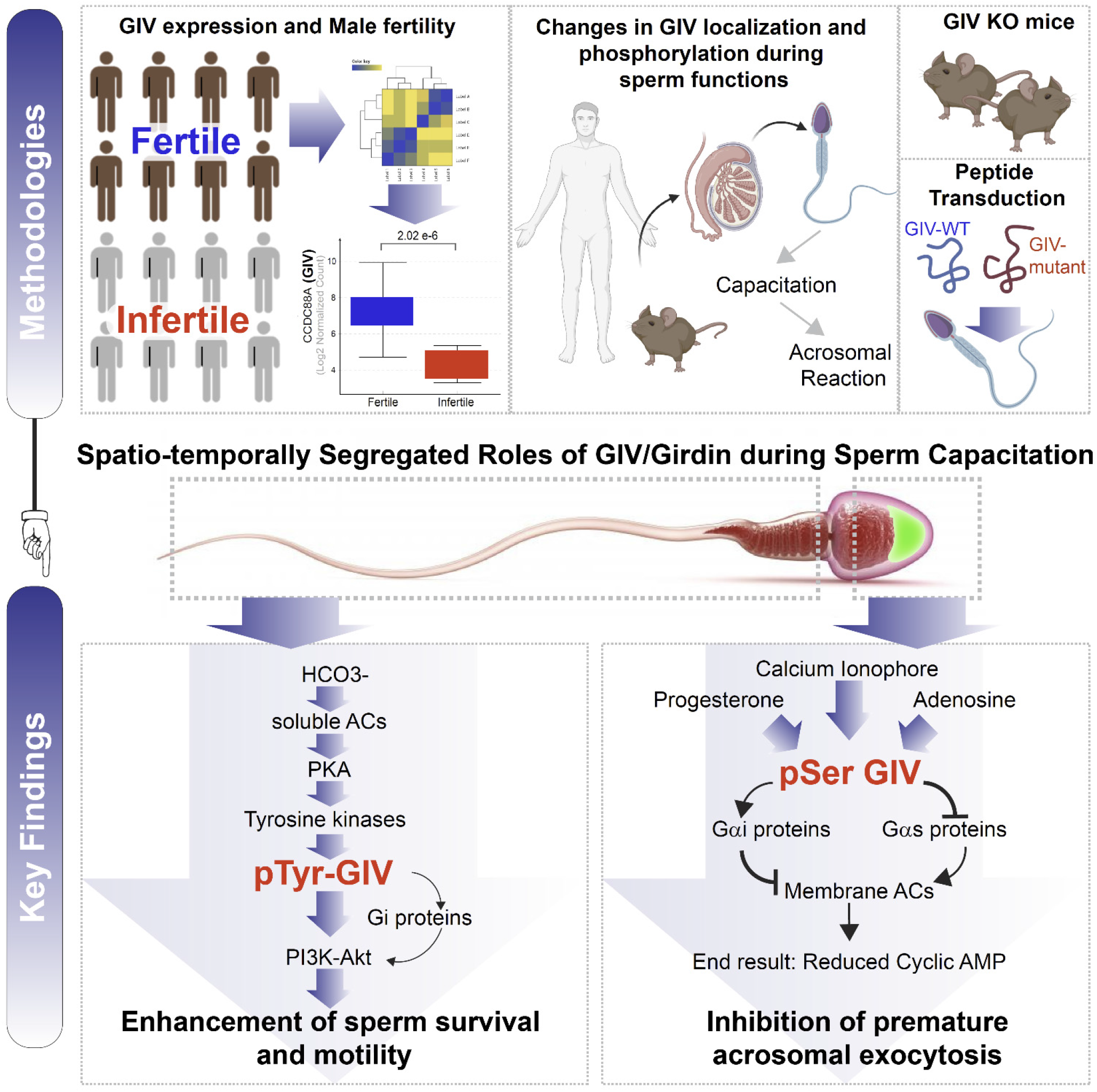

**HIGHLIGHTS:**

- GIV is highly expressed in spermatozoa, and is required for male fertility
- GIV is rapidly phosphoregulated during sperm capacitation
- It enhances tyrosine-based signals in sperm tail, enhances motility
- It suppresses cAMP in the sperm head, inhibits premature acrosome exocytosis

## INTRODUCTION

Mammalian sperm acquire their fertilizing potential after insemination, during the passage through the female reproductive tract. Two key consecutive processes are prerequisites for successful fertilization: (i) sperm must first undergo capacitation, a process that is characterized by progressive acquisition of hypermotility, change in membrane, and phosphorylation status, and (ii) they must later undergo acrosome reaction (AR), a process that is characterized by an exocytotic release of acrosomal enzymes to penetrate the zona pellucida of the egg^1,2^. Although capacitation is an important physiological pre-requisite before spermatozoa can fertilize the oocyte in every mammalian species studied, the molecular mechanisms and signal transduction pathways involved in this process are poorly understood. AR, on the other hand, is a time-dependent phenomenon that cannot take place prematurely or too late^3^. Premature spontaneous AR that occurs in the absence of proper stimuli (AR insufficiency) has been associated with idiopathic male infertility^4^.

Being transcriptionally and translationally silent, mature spermatozoa support capacitation and AR relying exclusively on post-translational events, e.g., increase in membrane fluidity, cholesterol efflux, ion fluxes resulting in alteration of sperm membrane potential, and an increased protein phosphorylation; the latter represents a very important aspect of capacitation^5^ (summarized in **==Figure 1-figure supplement 1A-B**). Despite these mechanistic insights into sperm capacitation, key gaps in knowledge persist. For example, although it is known that phosphotyrosine intermediates in the sperm tail culminate in the activation of the PI3K→ Akt signaling axis, and that such activation is vital for sperm hypermotility, how tyrosine phosphorylation leads to the activation of PI3K remains unknown^6,7^. Similarly, although it is known that Akt-dependent actin polymerization in the sperm tail requires both protein kinase A and protein tyrosine phosphorylation, the linker(s) between signaling and actin dynamics remains unidentified^7-9^. Finally, how cAMP surge during capacitation is restricted to the sperm tail, such that its levels remain low in the sperm head, and premature AR is avoided, remains a mystery.

Here we show that GIV (a.k.a., *GIRDers of actIN filament*, Girdin; gene: CCDC88A), a multimodular signal transducer, straddles both tyrosine-based and G protein→ cAMP signaling cascades during sperm capacitation. There are several known functions of GIV, all reversibly modulated by phosphorylation cascades, that make it an ideal candidate to fill some of the knowledge gaps: (i) As a substrate of multiple tyrosine kinases (RTKs and non-RTKs, alike) two phosphotyrosines within GIV’s C-terminus directly bind and activate Class 1 PI3Ks^10,11^; this makes GIV a point of convergence for multi-TK-dependent PI3K signaling. (ii) As a *bonafide* enhancer and a substrate of Akt^12^, GIV binds and depolymerizes actin and the only known substrate of Akt that links the PI3K→ Akt cascade to cytoskeletal remodeling^12^. (iii) As a guanine nucleotide-exchange modulator (GEM) for trimeric GTPases, GIV serves as a GEF for Gi^13^ and a GDI for Gs^14^ via the same evolutionarily conserved C-terminal motif. By activating the inhibitory Gi and inhibiting the stimulatory Gs proteins, GIV overall inhibits membrane adenylyl cyclase (mACs) and suppresses cellular cAMP^15^. ‘Free’ Gβγ that is released from the Gi/s trimers further enhances the PI3K→ Akt signals^13^. We show here how GIV orchestrates distinct spatiotemporally segregated signaling programs in sperm to support capacitation and concomitantly inhibit premature AR, thereby playing an essential role in male fertility.

## RESULTS AND DISCUSSION

### GIV is highly expressed in spermatocytes

At the time of its discovery in 2005, full length GIV protein was found to be most highly expressed in two organs: testis and brain (**Fig 1A**). Immunohistochemical studies curated by the Human Protein Atlas further confirm that GIV is most highly expressed in the testis (**Figure 1- figure supplement 2A**). Single cell sequencing (**Fig 1B; Figure 1- figure supplement 2B-E**) and IHC (**Fig 1C**) studies on human testis pinpoint sperm as the major cell type in the testis that expresses GIV mRNA and protein. We confirmed by confocal immunofluorescence on mouse testis that GIV is indeed expressed in the spermatozoa, and localizes predominantly to the acrosomal cap, as determined by colocalization with the mouse acrosomal matrix protein, sp56^16^ (tGIV; **Fig 1D**). As expected, a tyrosine phosphorylated pool of GIV (pYGIV), however, localized mostly to the plasma membrane (PM) (**Fig 1E**). Both antibodies detected the endogenous GIV protein in testicular lysates at the expected size of ∼ 220 kDa (**Fig 1F**). We also noted that GIV consistently and predominantly localizes to the acrosome, as it matures from a rudimentary vesicle into a vesicular cap during sperm maturation (**Fig 1G**). GIV’s localization to the acrosome, which is derived from the Golgi apparatus^17^, is in keeping with GIV’s predominant localization to the Golgi and Golgi-associated transport vesicles in diverse cell types^18,19^. Taken together, we conclude that GIV is highly expressed in sperm, and may be important for sperm functions.

**Figure 1.**
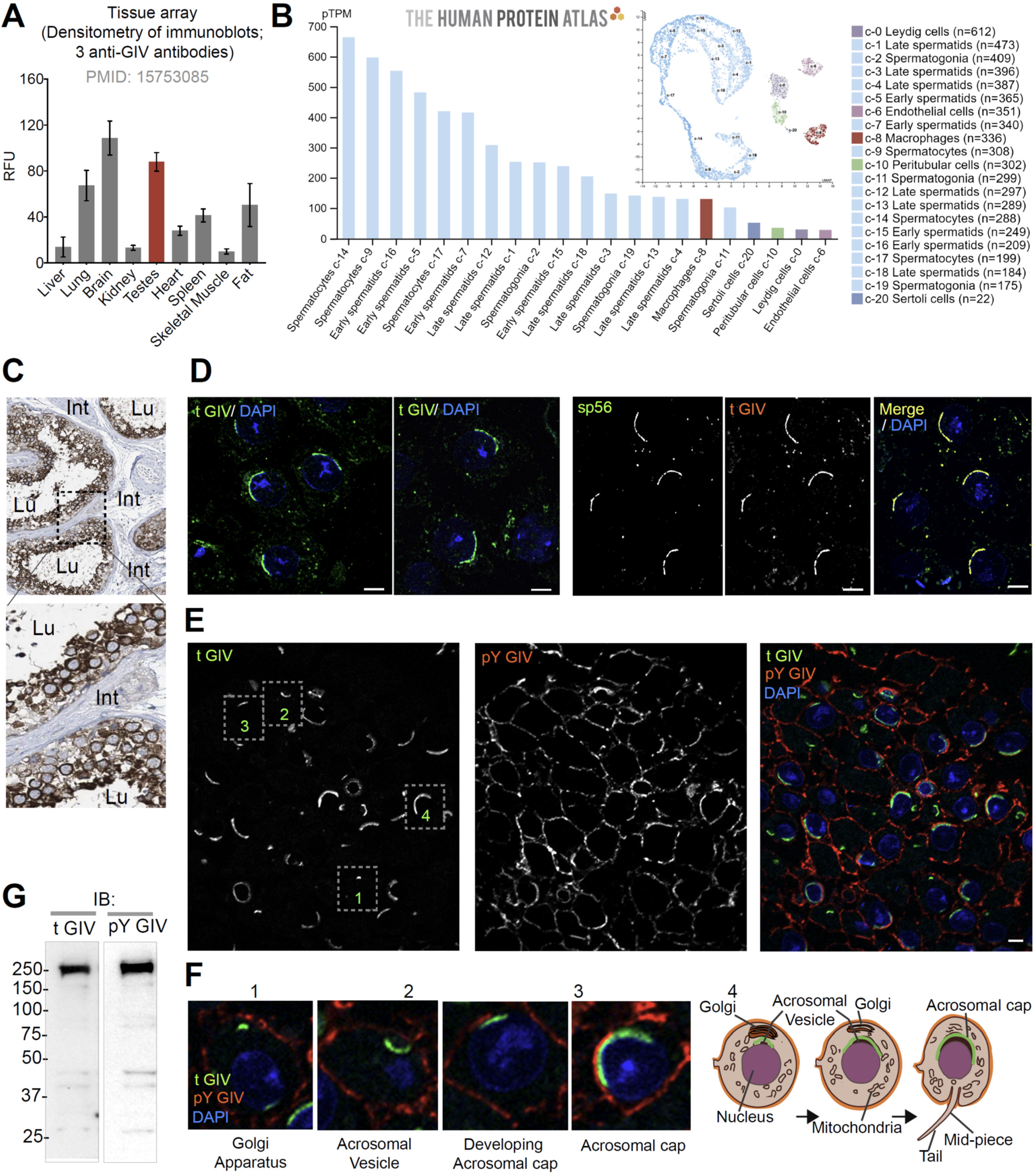
GIV (CCDC88A) is highly expressed in spermatocytes in testis and localizes to the acrosomal cap. **A**. Bar graph displays the relative fluorescence unit (RFU) of endogenous full length GIV protein in immunoblots of organ lysates published previously using three independent anti-GIV antibodies raised against different epitopes of GIV^64^. **B**. RNA expression in the single cell type clusters identified in the human testis visualized by a UMAP plot (inset) and a bar plot. The bar plot shows RNA expression (pTPM) in each cell type cluster. UMAP PLOT visualizes the cells in each cluster, where each dot corresponds to a cell. **C**. Representative images from human testis IHC studies curated in the HPA. Int = interstitium; Lu = lumen of seminiferous tubules. **D**. Cryosections of mouse testis (8 wk old, C57BL/6) were stained for either total GIV (tGIV; green) and DAPI (blue, nucleus) alone, or co-stained with tGIV and the sperm acrosomal matrix protein zona pellucida 3 receptor (ZP3R, formerly called sp56; red) and analyzed by confocal immunofluorescence. Representative images are displayed. Scale bar = 10 µm. **E-F**. Cryosections of mouse testis tissue analyzed for total (t) GIV (green), pY GIV (red) and DAPI (blue, nucleus). Representative images are showing in panel E; Scale bar = 10 µm. Insets in panel F are magnified and displayed in panel E, left. Schematics in panel G, right display various localization of GIV observed during the process of maturation of the Golgi into acrosomal cap. **G**. Immunoblots on mouse testis lysates with the same tGIV and pY GIV antibodies.

### Transcripts of GIV are reduced in infertile men

Previously, in a publicly available patent (WO2017024311A1), the GIV gene (CCDC88A) was identified as one among a panel of genes whose altered expression due to DNA methylation may help diagnose male fertility status and/or the quality of the embryo^20^. We asked if the abundance of GIV transcripts in testis or sperm may be altered in infertile men. To this end, we curated all publicly available transcriptomic datasets from the NCBI GEO portal and analyzed them for differences in the abundance of CCDC88A transcripts across the annotated (in)fertility phenotypes (**Fig 2A**). CCDC88A transcripts were significantly and consistently downregulated in infertile men across all independent datasets analyzed (**Fig 2B-E**), regardless of whether the samples used for transcriptomic studies were testis or sperm. Finally, in a study that segregated sub-fertile from fertile men using commonly used clinical parameters for semen quality, we found that sperm motility, but not concentration or morphology was the key parameter (**Fig 2F**); reduced motility in sub-fertile men was associated also with reduced levels of GIV transcript. These results indicate that reduced GIV expression in testis and sperm is associated with clinically determined male infertility. Given the heterogeneous nature of the datasets (i.e., diagnosed cause of infertility, ranging from genetic syndromes with developmental or hormonal defects to post-chemotherapy to idiopathic), reduced GIV expression could be considered as a shared common molecular phenotype among infertile men.

**Figure 2.**
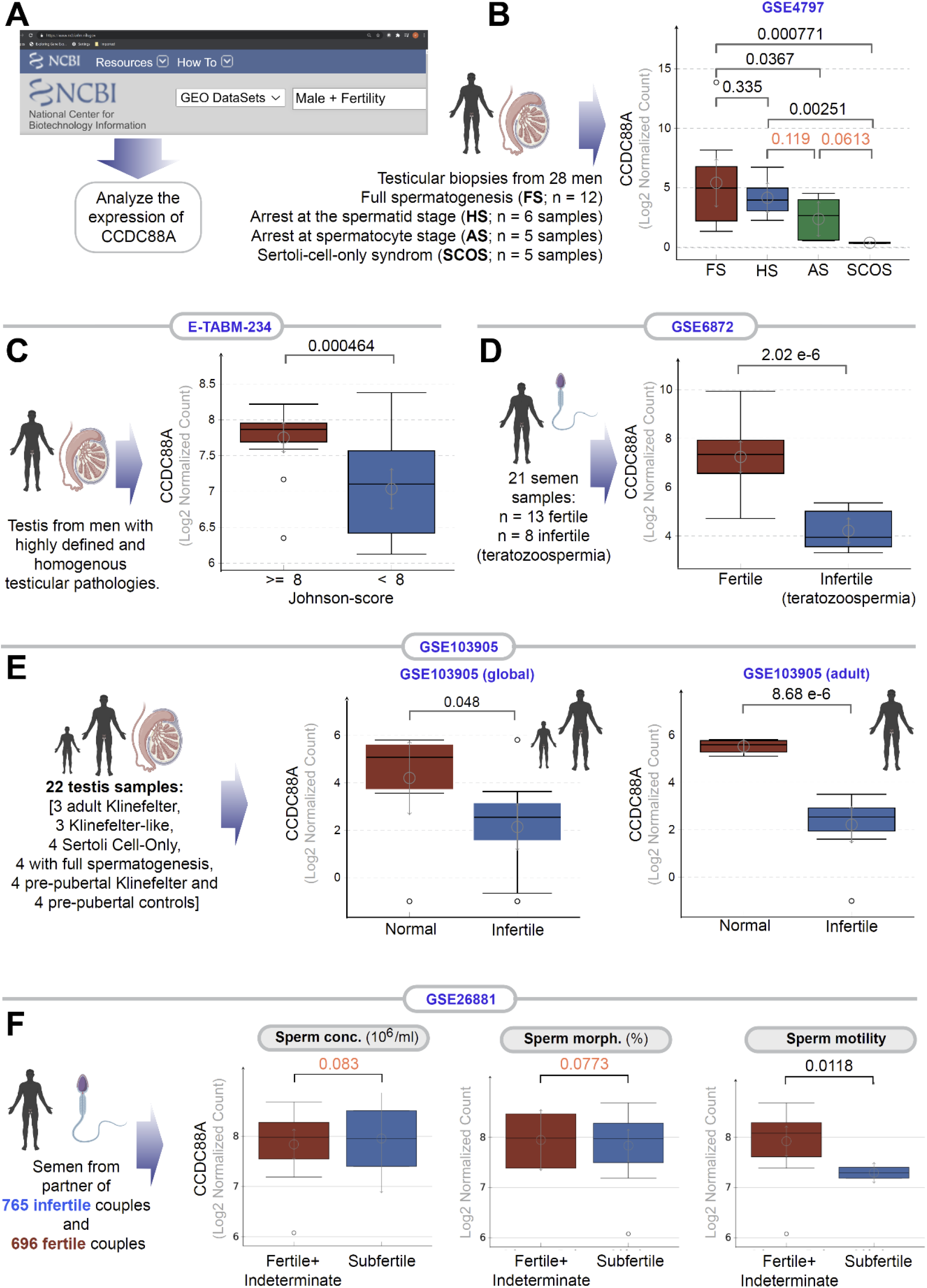
Transcripts of CCDC88A (GIV) is downregulated in infertile male testis and semen. **A**. Schematic displays the approach used to search NCBI GEO database for testis and sperm transcriptomic datasets suitable to study correlations between the abundance of CCDC88A transcripts and male fertility. **B-E**. Whisker plots show the relative abundance of CCDC88A (expressed as Log2 normalized transcript count) in sperm or testis samples (as annotated using schematics) in samples annotated with fertility status, or syndromes associated with infertility. **F**. Whisker plots shows the relative abundance of CCDC88A transcripts in sperms classified as sub-fertile or not based on three properties of sperm assessed using a modified WHO criterion published by Guzick et al^65^ (see *Methods*).

### GIV is rapidly tyrosine phosphorylated during capacitation

Mature sperm, by virtue of being transcriptionally and translationally inactive, rely entirely upon rapid post-translational modifications to regulate all pre-zygotic processes. Because GIV is a multimodular signal transducer that straddles both tyrosine based and G protein signaling pathways^21,22^, we sought to investigate how GIV’s functions are altered during sperm capacitation. Because PI3K-Akt signals downstream of tyrosine kinases is a critical pathway for actin remodeling in the sperm flagellum and for hypermotility^6-9^, and GIV serves as a point of convergence for multi-TK-dependent PI3K signaling^10,11^, we first asked if GIV is indeed tyrosine phosphorylated in human and mouse sperm during capacitation. Using the sperm swim-up assay, we first confirmed that in human ejaculates, low motile sperm have just as much total GIV as their highly motile counterparts, but by contrast, tyrosine phosphorylated GIV was significantly elevated in the latter (compare tGIV and pYGIV, lanes 1-2 in immunoblots; **Fig 3A**). As a positive control, we simultaneously analyzed the same samples by dual color immunoblotting with an antibody that detects pan-tyrosine phosphoproteins. As expected^23-26^, the highly motile sperms have higher tyrosine phosphorylation (pan-pY; **Fig 3A**). pYGIV and pan-pY signals co-migrated in the SDS page gel, indicating that GIV is one of the tyrosine phosphorylated proteins in high motile sperms. The pan-pY and pYGIV signals were found to further increase in capacitated sperms, maximally by 4 h, without any change in total GIV (lanes 3-4; **Fig 3A**). Such phosphorylation was dependent on the activity of protein kinase A (PKA) because pre-treatment of sperm with the PKA inhibitor H89 virtually abolished both pan pY and pYGIV (**Fig 3B**); these findings are in keeping with the fact that PKA activity is essential for tyrosine phosphorylation cascades during capacitation^27,28^. Immunofluorescence studies on human sperm confirmed that pan-pY and pYGIV signals colocalized in the mid-piece and tails of high-motile sperm (**Fig 3C**) where they were significantly induced upon capacitation (**Fig 3D**). Findings in human sperm were mirrored in murine sperm (**Fig 3E-F**), with some notable differences in temporal-spatial dynamics. For example, pY/pYGIV of murine sperms are induced more rapidly and transient. During murine sperm capacitation pYGIV is induced in 30 min and then reduced in 120min (**Fig 3F**) and was not as restricted to the sperm tail and mid-piece as in humans (compare sperm head regions in **3D** with **3F**). Although full length GIV (∼250 kDa expected size) could be detected in murine sperm (**Fig 3F**), we often detected numerous breakdown products, presumably proteolytic in nature, in both murine and human sperm lysates (**Fig 3A-B, F**). Regardless of the size of the breakdown products, total tGIV, pYGIV and pan-pY co-migrated in the gels at the same size, suggesting that GIV may be one of the major phosphotyrosine proteins in capacitating sperm. We conclude that GIV is a major phosphotyrosine substrate in sperm tail during capacitation and that its phosphoactivation requires upstream activation of PKA. Our findings suggest that this PKA→ TK→ pYGIV axis may enhance PI3K-Akt signals and sperm motility.

**Figure 3.**
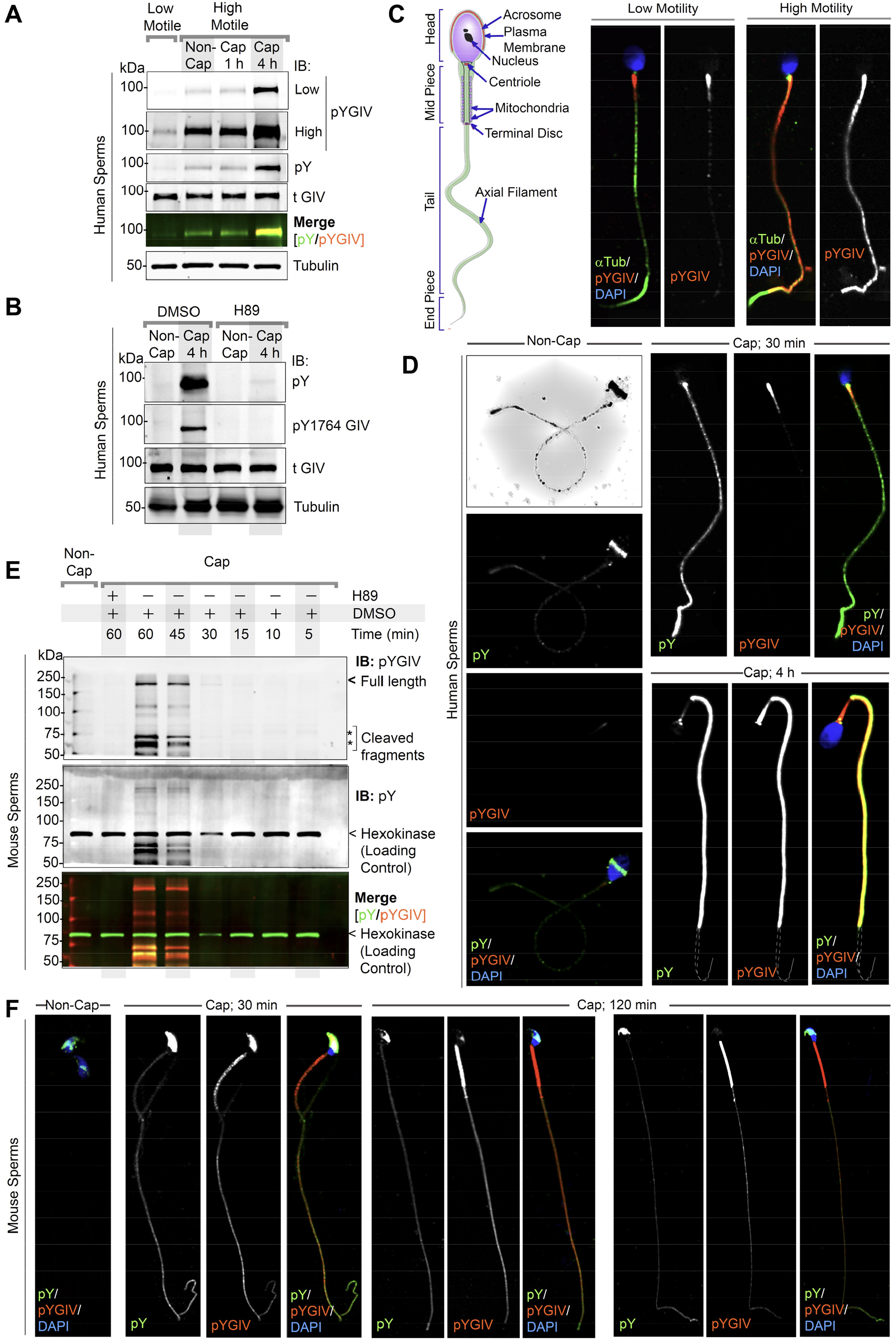
GIV localizes to the head and tail of human and murine sperms, is rapidly tyrosine phosphorylated during capacitation. **A**. Freshly ejaculated human sperm were segregated into low motile and high motile populations using ‘swim-up’ technique (see Methods) and subsequently capacitated in vitro for 1 or 4 h prior to whole cell lysis. Equal aliquots of lysates were analyzed by immunoblotting for total (t) GIV, pan pY, pY1764 GIV (pYGIV) and β-Tubulin (loading control) using LI-COR Odyssey. **B**. Whole cell lysates of human sperms capacitated with or without preincubation with H89 (PKA inhibitor) or DMSO control were analyzed as in A. **C-D**. Human sperm with low vs high motility (C), were capacitated or not (D), fixed and co-stained for total and pY GIV (tGIV; pY GIV), tubulin and DAPI. Scale bar = 10 µm. **E**. Immunoblots of equal aliquots of whole cell lysates of mouse sperm capacitated with (+) or without (-) pre-treatment with PKA inhibitor (H89) or vehicle (DMSO) control. Hexokinase is used as a loading control. **F**. Non-capacitated (non-cap) or capacitated mouse sperm were fixed and stained as in D and analyzed by confocal microscopy. Scale bar = 10 µm.

### The G protein modulatory function of GIV is dynamically phosphoregulated during capacitation

Next, we asked how the G protein modulatory function of GIV is regulated during capacitation. The evolutionarily conserved C-terminal GEM motif in GIV that enables it to both activate Gi^13^ and inhibit Gs^14^ is phosphoregulated by two Ser/Thr kinases, Cyclin-dependent-like kinase 5 (CDK5)^29^ and Protein kinase C ⊖ (PKC⊖)^30^ (summarized in **Fig d4A**). Phosphorylation at S1674 induces GIV’s ability to activate Gi by 2.5-3-fold, whereas phosphorylation at S1689 inhibits GIV’s ability to activate Gi; neither phosphoevent impacts GIV’s ability to bind and inhibit Gs. By activating the inhibitory Gi and inhibiting the stimulatory Gs proteins, GIV overall inhibits membrane adenylyl cyclases (mACs) and suppresses production of cellular cAMP^15^. Because post-translational protein modification is the predominant way mature sperm rapidly respond to environmental cues, we used two previously validated phosphosite-specific antibodies^29,30^ that detect pS1674-GIV and pS1689-GIV. We found that in mouse (**Fig 4B**; left) and human (**Fig 4C**; left) sperm, HCO3^-^-induced capacitation induced the levels of phosphorylation at the activation site pS1674 in the sperm tails of both species, with two notable inter-species differences—(i) In murine sperm the acrosomal cap showed phosphorylation at baseline with no further increase upon capacitation, and (ii) In human sperm the mid-piece region showed phosphorylation at baseline with no further increase upon capacitation. Unlike pS1674, distribution/intensity of phosphorylation at the inhibitory pS1689 site was observed at baseline in the head, mid-piece, and tail of the murine sperm (**Fig 4B**, right), and the mid-piece and tail in human sperm (**Fig 4C**, right) and did not change during capacitation.

**Figure 4.**
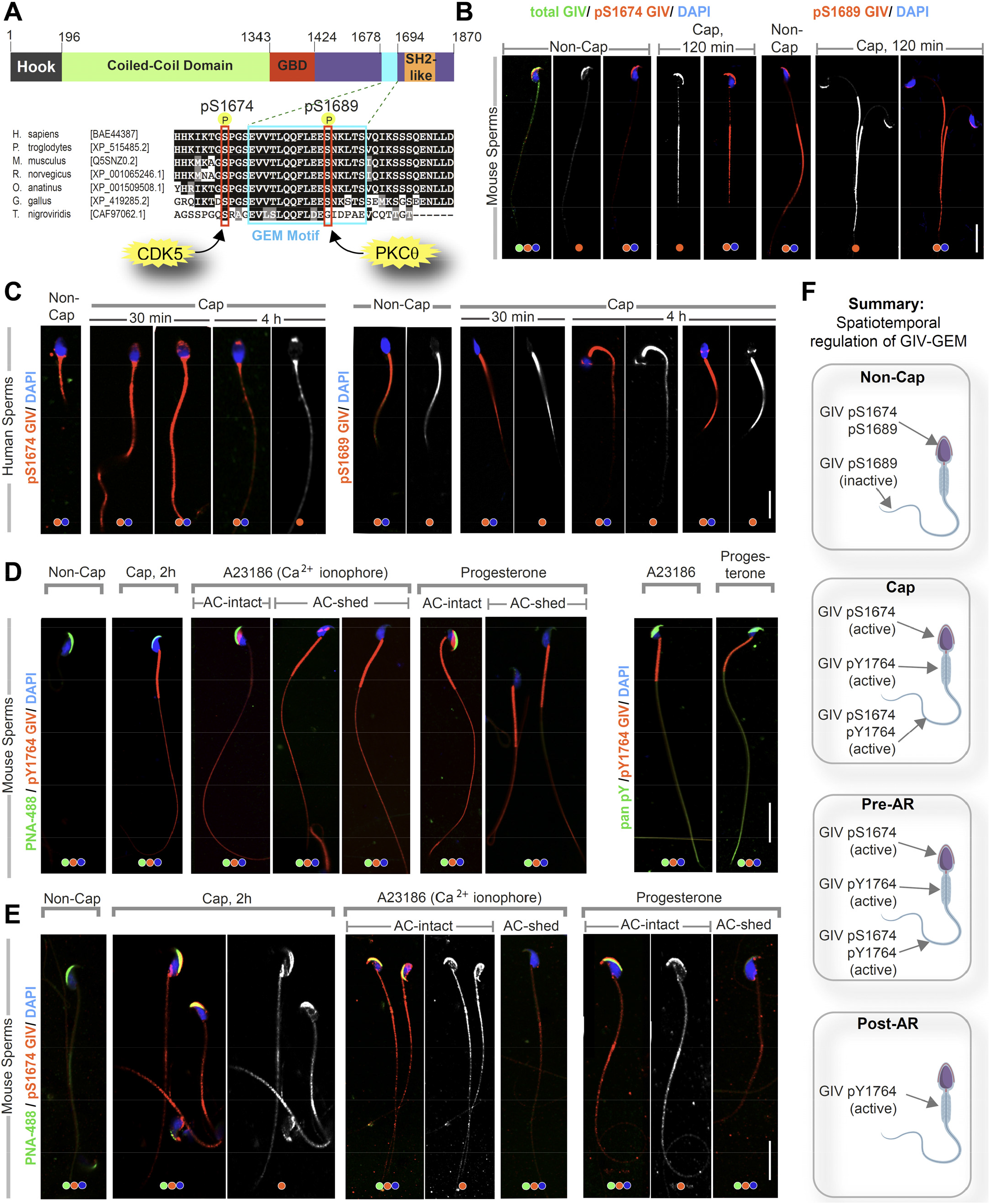
GIV’s GEM function is dynamically phosphoregulated during capacitation and acrosomal reaction in a spatiotemporally segregated manner. A. Schematic shows the domain map of GIV (top) and the evolutionarily conserved GEM motif within its C terminus. A functional GEM motif is required for GIV to bind and activate Gαi as well as bind and inhibit Gαs^14^. Important phosphoserine modifications that regulate GIV’s GEM motif and the corresponding target kinases are highlighted. **B-C**. Non-capacitated and capacitated mouse (B) and human (C) sperm were fixed and analyzed for the phosphoserine modifications highlighted in A. **D-E**. Mouse sperm with/without capacitation followed by treatment with either Ca^2+^ ionophore or progesterone to trigger acrosomal reaction were fixed and co-stained for peanut agglutinin (PNA-488; green, an acrosomal marker) and either pYGIV (D) or pSerGIV (E) and DAPI. Representative images are shown. Scale bar = 10 µm. **F**. Schematic summarizes the spatially segregated phosphomodifications on GIV before and after capacitation and acrosomal reaction (AR) in various parts of the sperm. (i) inhibitory phosphorylation at pS1689 on GIV is seen in both head and tail prior to capacitation (**F**, top); (ii) activating phosphorylation at S1674 on GIV is seen in the sperm head and tail, whereas pYGIV is predominantly seen in the mid-piece and the tail regions upon capacitation (post-cap; **F**) as well as during acrosomal reaction before the acrosome is shed (pre-AR; **F**); (iii) after the acrosome is shed, pYGIV is the only phospho-GIV that is detected, and predominantly in the mid-piece (post-AR; **F**).

We next repeated the studies with the sequential addition of HCO3^-^ (for 2 h) followed by two other stimuli that are commonly used to trigger the acrosome reaction, the calcium ionophore, A23186^31^ and the reproductive hormone, progesterone^32^. To monitor the phosphomodifications in GIV and their temporal relationship with acrosome exocytosis, we co-stained the sperm with peanut agglutinin (PNA) and Ser/Tyr-GIV. PNA binds specifically to galactose residues on the outer acrosomal membrane, and its disappearance is a widely accepted method of monitoring acrosome exocytosis^33,34^. pYGIV was induced predominantly in the mid-piece and tail during capacitation (cap 2h; **Fig 4D**) as seen before (**Fig 3F**) but also in the sperm head in the presence of A23187 and progesterone (**Fig 4D**). The localization of pYGIV in sperm head was seen only when the acrosomes were intact and lost in those where the acrosome was shed (compare AC-intact vs -shed; **Fig 4D**). Similarly, phosphorylation at the activation site pS1674 was detected in sperm heads in the presence of A23187 and progesterone, but exclusively when the acrosomes remained intact (compare AC-intact vs -shed; **Fig 4E**).

Taken together, the predominant findings can be summarized as follows (see legend **Fig 4F**): GIV-GEM is inactive at baseline and activated upon capacitation. It remains active in both head and tail regions of capacitated sperm until the moment the acrosome is shed. Capacitation is also associated with robust tyrosine phosphorylation of GIV in the sperm tail and mid-piece throughout the process of acrosomal reaction.

### GIV is required for male fertility

To determine if GIV is required for male fertility, we next co-housed female mice with conditional GIV-/-male mice (GIV-KO; generated using Tamoxifen in Girdin fl/fl-UBC-Cre-ERT2 mice) or control littermates (WT; Girdin fl/fl mice) (see methods; legend **Fig 5A**) and analyzed diverse readouts. GIV knockdown was confirmed by genotyping tail tips (**Fig 5B**) and by assessing GIV mRNA (**Fig 5C**) and protein (**Fig 5D**) in the testis. We noted a significant reduction of cumulative probability of pregnancy (100% vs. 55% rate for WT and KO group, respectively within 40 days after co-housing; **Fig 5E**) and average litter size (**Fig 5F**) in GIV-KO mice. Surprisingly, both WT and GIV-KO mice had similar sperm counts (**Fig 5G**), testes sizes and weights (**Figure 5- figure supplement 1**). We confirmed by IHC that GIV was predominantly expressed in sperm in the testis of WT mice and that it was effectively depleted in GIV-KO mice (**Fig 5H**). RNA seq of the testis followed by unsupervised clustering showed that GIV-KO testis differentially expressed only a handful of transcripts compared to WT testis (**Fig 5I**). The predominantly upregulated genes mapped to the ‘aberrant activation of PI3K/Akt signaling’ pathway (**Fig 5J**). This was largely attributable to *Esr1* (highlighted in red; **Fig 5I**); polymorphisms of this gene are known to predispose to male fertility^35,36^ and its induction represents a negative feedback event resulting in the setting of inhibition of PI3K signaling^37^. The predominantly downregulated genes mapped to the IL12 pathway (**Fig 5K**), which is consistent with prior studies in men showing that IL12 may be important for male fertility and that its dysregulation may reflect infertility^38,39^. Notably, both pathways reflect changes that are largely contributed by non-sperm cells in the testis; Esr1 is expressed exclusively in the Leydig cells in mouse testis^40,41^ and IL12 is largely expressed by endothelial cells, peritubular cells and macrophages^42^.

**Figure 5.**
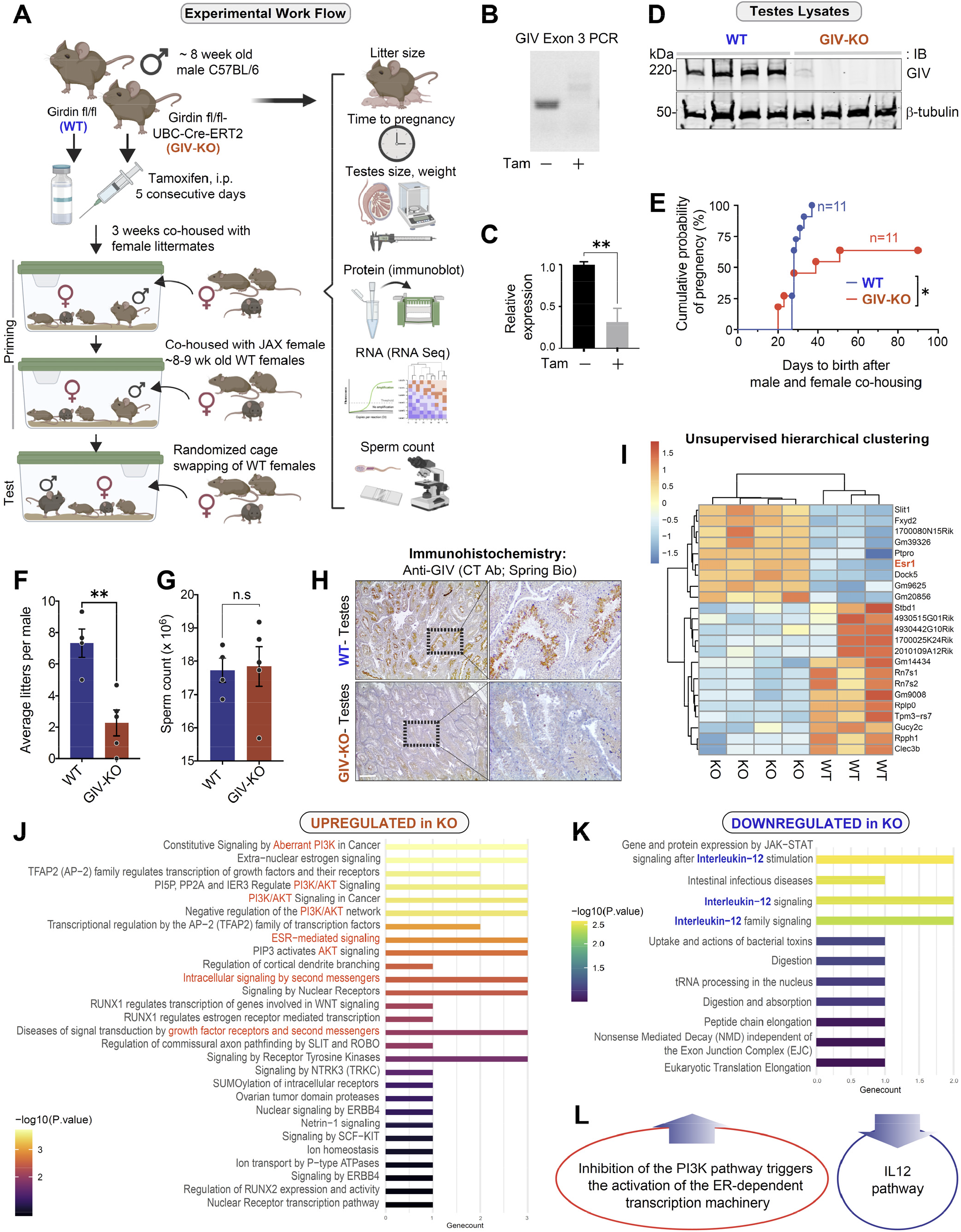
GIV is required for fertility in male mice. **A**. Schematic showing the workflow for fertility studies in conditional GIV-KO mice. After intraperitoneal injection of Tamoxifen, male mice were first primed in 2 phases— first by co-housing with female littermates x 3 weeks, and subsequently by co-housing with female mice from Jackson laboratory (JAX) while the females acclimatized to the animal facility. The final ‘test’ group consisted of Tamoxifen-injected WT and GIV-KO male mice randomly assigned to and co-housed with 3 female mice from JAX, each with proven ability to get pregnant. **B-D**. Confirmation of GIV-KO in the mice after tamoxifen injection by genotyping (B), qPCR of testis tissues (C) and immunoblotting of testis lysates (D). **E**. Kaplan-Meier plot showing the cumulative probability of pregnancy (expressed as %) in the females co-housed with either WT or GIV-KO males. **F-G**. Bar graphs showing the average litter size (F) and sperm count (G) in WT and GIV-KO males. **H**. IHC staining on mouse testis. Scale bar = 200 µm. **I**. Unsupervised clustering of WT and KO testis samples based on gene expression. DEGs that were up or downregulated in KO are annotated on the right side. **J-K**. Reactome pathway analyses showing the pathways that are UP or DOWN-regulated in KO testis. **L**. Summary of the most prominent conclusions from RNA seq dataset.

These findings demonstrate that GIV is required for male fertility and suggests that the role of GIV and its various phosphomodifications we observe in sperm is largely post-transcriptional and post-translational in nature.

### GIV’s GEM function facilitates hypermotility and survival during sperm capacitation

Next, we assessed the role of GIV during sperm capacitation using a previously validated approach, i.e., exogenous addition of cell-permeable His-tagged GIV-derived ∼210 aa long peptides^43^; these peptides either have an intact functional GEM motif (WT peptides) or, as negative control, a well-characterized F1685A (FA) mutant of the same motif which lacks such activity^13,44^ (**Fig 6A**; top). By anti-His staining followed by flow cytometry, we confirmed that TAT-His-GIV peptides were indeed taken up as we could detect uptake only when staining was conducted under permeabilized conditions (**Fig 6A**; bottom). Peptide uptake was efficient, varying within the range of ∼80-90% (**Fig 6A**; bottom). Immunofluorescence studies confirmed that uptake was seen in all segments of the sperm (**Fig 6B**). The peptides were detected and functional (i.e., retained their ability to bind Gαi) at 1 and 6 h post-uptake, as determined using lysates of peptide-transduced sperm as source of GIV in pulldown assays with recombinant GDP-loaded GST-tagged G protein, Gαi3 (**Fig 6C**).

**Figure 6.**
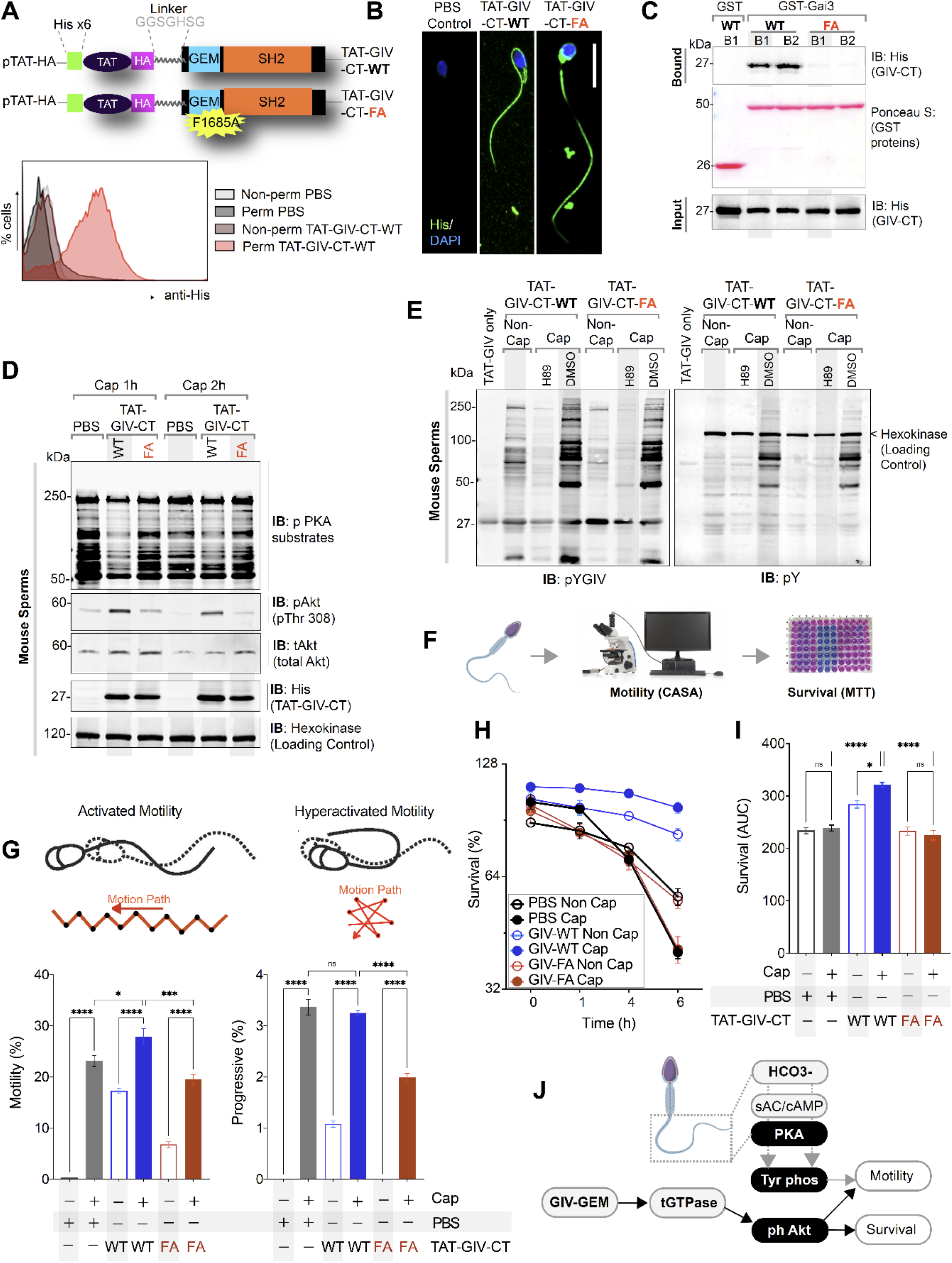
GIV’s GEM function is required for sperm motility and survival during capacitation. **A-B**. Schematic (A, top) of cell permeant His-TAT-GIV-CT wild type (WT) and GEM-deficient mutant (F1685A; FA) peptides used in this work. Immunofluorescence images (B) representative of sperms after treatment with cell-permeant TAT-GIV-CT peptides and stained with anti-His antibody and DAPI. Scale bar = 15 µm. Histograms (A, bottom) from flow cytometry studies conducted with or without permeabilization confirm the uptake of His-TAT peptides in sperm. **C**. Immunoblots of GST pulldown assays testing the ability of GDP-loaded GST-Gαi3 to bind TAT-GIVCT peptides from lysates of sperms at 1 h (B1) and 6 h (B2) after transduction. **D-E**. Immunoblots of lysates of TAT-GIVCT transduced sperms at the indicated time points after capacitation analyzed for phospho-PKA substrates (D), phospho(p) and total (t) Akt (D), pYGIV (E, left), pan-pY (E, right) and hexokinase (loading control, D). **F-I**. Schematic in F summarizes workflow in assessing motility and survival of sperms during capacitation. Bar graphs in G display the relative % of motile and progressively motile population of sperms. Line graphs in H show survival of sperms as determined by MTT assay; bar graphs in I show the AUC of the line graphs in H. **J**. Schematic summarizes the conclusions of how GIV’s GEM function impacts sperm phenotypes during capacitation.

Next, we analyzed phosphoproteins in TAT-GIV-transduced sperm undergoing *in vitro* capacitation by immunoblotting. Although PKA activation (**Fig 6D**) and pan-Y or pYGIV phosphorylation (**Fig 6E**) were relatively similar between WT and FA transduced sperm, phosphorylation of Akt differed; TAT-GIV-WT, but not FA induced phosphorylation of Akt (**Fig 6D**). This finding is consistent with the established role of GIV-GEM in the activation of the PI3K→ Akt pathway via the activation of Gi and the release of ‘free’ Gβγ^13^. Because Akt phosphorylation has been implicated in sperm hypermotility and survival during capacitation^45,46^, we performed computer-assisted sperm analysis (CASA) and MTT assays, respectively (**Fig 6F**). Consistent with the patterns of Akt phosphorylation, WT, but not FA peptide-transduced sperm showed greater overall motility as well as hypermotility (**Fig 6G**) and greater viability (**Fig 6H-I**).

These findings indicate that GIV’s GEM function may be dispensable for the PKA→ TK→ tyrosine phosphorylation pathway, but is required for Akt activation, sperm motility and survival during capacitation (**Fig 6J**).

### GIV’s GEM function suppresses cAMP and acrosomal reaction

Prior studies have underscored the importance of membrane Adenylyl Cyclases (mACs) and their role in the regulation of cAMP and acrosome exocytosis in sperm (summarized in **Figure 1- figure supplement 1A**). Membrane ACs are localized most abundantly in the head (**Figure 1- figure supplement 1A**), and their activation by Gs or inhibition by Gi is known to finetune cAMP surge in that location, spatiotemporally segregated and independent of the sAC-dependent cAMP surge in the sperm tail (**Figure 1- figure supplement 1B**). Upon approaching the zona pellucida of an egg, a timely surge in cAMP in sperm head is required for the downstream activation of effectors PKA^47^ and the exchange proteins directly activated by cAMP (EPAC)^48,49^, which in turn coordinate the rapid cytoskeletal remodeling and membrane trafficking events that lead to acrosome exocytosis. As an activator of Gi and an inhibitor of Gs^14^ using the same conserved GEM motif (**Fig 7A**) GIV is known to tonically and robustly suppresses cAMP^15,50^, and by that token, it is expected to inhibit the cAMP surge. Because GIV-GEM was activated upon capacitation and remained active until the acrosome was shed (**Fig 4F**), we hypothesized that GIV’s GEM function may be required for the prevention of a premature cAMP surge in the sperm head, and hence, premature acrosome exocytosis. We first confirmed that cAMP is modulated by a variety of stimuli targeting Gi-(adenosine) and Gs-coupled (progesterone) GPCRs (**Figure 7- figure supplement 1A**), consistent with what has been observed before^51,52^. When the same studies were carried out on TAT-GIV peptide transduced sperm,, the expected degree of cAMP induction were observed once again (**Fig 7B**), but TAT-GIV-WT, but not the GEM-deficient FA mutant peptides could significantly suppress the degree of cAMP surge across all stimuli tested (**Fig 7C; Figure 7- figure supplement 1B**).

**Figure 7.**
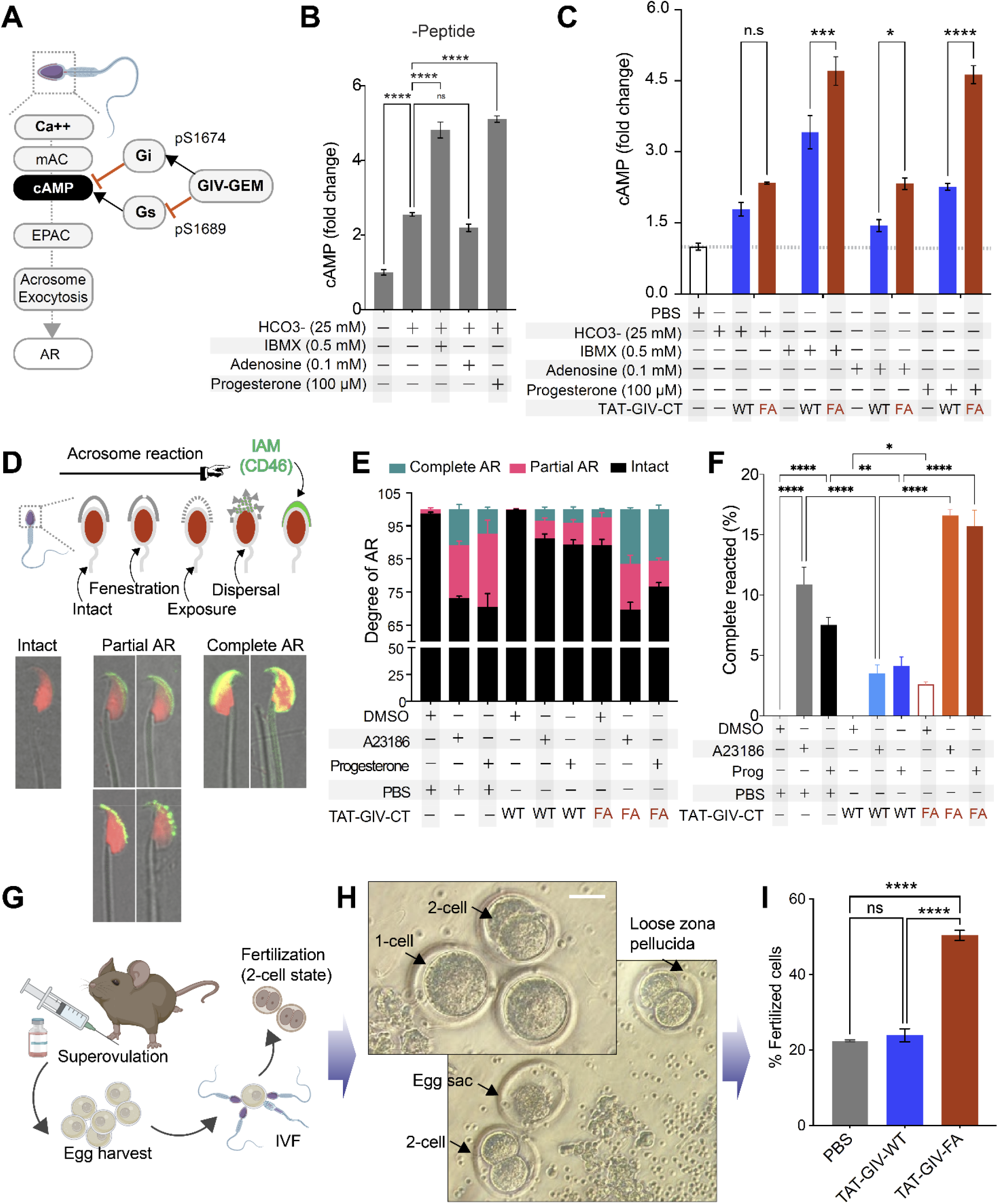
GIV’s GEM function inhibits acrosomal reaction. **A**. Schematic summarizes the current knowledge of how Ca^2+^ and cAMP signaling regulates acrosome exocytosis during acrosomal reaction (AR) and how GIV’s ability to modulate cAMP via both Gαi/s is hypothesized to impact AR. **B**. Bar graphs display the fold change in cAMP in mouse sperms treated with various stimuli in the presence of DMSO. **C**. Bar graphs display the fold change in cAMP in TAT-GIVCT transduced mouse sperms exposed to various stimuli. Dotted horizontal line represents cAMP concentration in PBS-treated samples, to which all other values were normalized. **D**. Schematic on top summarizes the assay used to quantify progressive changes in acrosome membrane during AR. Images in the bottom panel are representative of acrosome-intact, partial AR and complete AR stages. **E-F**. Stacked bar graphs in E display the proportion of sperms in each indicated condition that are either in partial or complete AR or with intact acrosomes. Bar graphs in F display just the relative proportion of sperms in E that have complete AR. **G-I**. Schematic in G displays the workflow used for *in vitro* fertilization (IVF) assays in H-I. Representative images in H display the 2-cell stage, which is quantified as % of total eggs in the assay and displayed as bar graphs in I as an indication of successful fertilization.

Next we assessed the effect of GIV-GEM on acrosome exocytosis under the same conditions using a highly sensitive immunofluorescence-based assay that monitors the progressive exposure during acrosomal reaction of the inner acrosomal membrane protein, CD46 (a.k.a^53,54^ membrane cofactor protein, MCP), (**Fig 7D**). TAT-GIV-WT, but not TAT-GIV-FA-transduced sperm had more intact acrosomes (**Fig 7E**) and fewer completely reacted acrosomes (**Fig 7F**), indicating that acrosomal reaction in response to both A23186 and progesterone was suppressed by TAT-GIV-WT, but not the GEM-deficient FA mutant.

These findings demonstrate that GIV is sufficient to inhibit cAMP surge and acrosome reaction, and that these functions require a functional GEM module. Taken together with the temporal nature of the GIV-GEM activity (see **Fig 4F**), our findings also suggest that GIV-GEM may inhibit premature cAMP surge and acrosome shedding. Because these premature events may compromise fertilization only in the *in vivo* setting where sperm is required to remain in capacitated state while maintaining intact acrosomes for prolonged periods of time within the female reproductive tract before encountering the egg, we hypothesized that GIV’s function may be bypassed in the setting of *in vitro* fertilization (IVF; **Fig 7G**). We found this indeed to be the case because TAT-GIV-WT peptide-transduced sperm successfully fertilized the eggs *in vitro* to a similar extent as PBS control (**Fig 7H-I**). The GEM-deficient FA mutant-transduced sperm, which had higher surges in cAMP (**Fig 7C**) and a higher proportion of completely reacted acrosomes (**Fig 7F**) showed a ∼2-fold increase in fertility.

Taken together, these findings indicate that GIV-GEM inhibits cAMP surge and acrosome reaction to primarily prevent both events from occurring prematurely *in vivo* until in the presence of an egg for successful fertilization.

## SUMMARY AND CONCLUSIONS

The major discovery we report here is a role of GPCR-independent (hence, non-canonical) G protein signaling in the sperm that is mediated by GIV/Girdin. Expressed most abundantly in the testis, and primarily, in sperm, GIV is required for male fertility, and low GIV transcripts in men was invariable associated with infertility. Mechanistically, we show that GIV is rapidly phosphomodulated on key Tyrosine and Serine residues in a manner segregated in space and time in various segments of the sperm (head, mid-piece and tail) during capacitation and acrosomal reaction. These specific phosphomodifications, which are known to regulate GIV’s interactions with other key proteins (PI3K, Gαi/s proteins, etc) and its functions as an effector of multiple TKs, as a cytoskeletal remodeler, and as a signal transducer regulate key sperm phenotypes in at least two sperm compartments (summarized in **Fig 8**):

**Figure 8.**
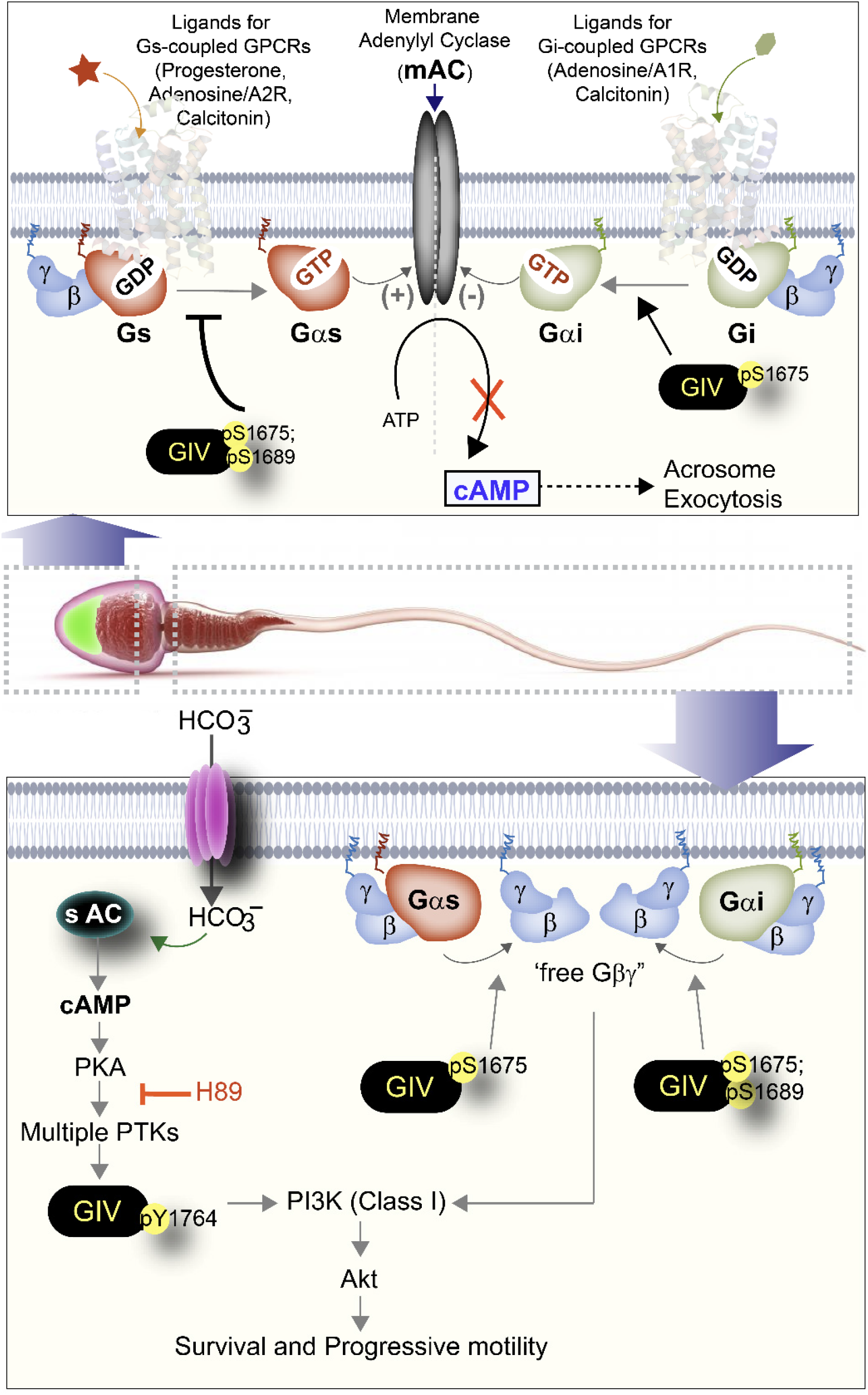
Summary and working model: Spatiotemporally segregated roles of GIV/Girdin during sperm capacitation. Schematic summarizes the key findings in this work and places them in the context of existing literature. GIV is likely to primarily function during capacitation of sperm, during which it fulfills two key roles as a signal transducer in a spatiotemporally segregated manner. The first role (Right, top) is in the head of the sperm, where GIV’s GEM motif inhibits the AC→ cAMP pathway and prevents acrosomal reaction. The second role (Right, bottom) is in the mid-piece and tail region of the sperm, which involves tyrosine phosphorylation of GIV, which happens downstream of PKA activation. Such phosphorylation is rapidly induced during capacitation. In addition, GIV’s GEM motif is activated, and is required for the enhancement of PI3K/Akt signals, enhanced motility and survival of sperms during capacitation.

In the sperm head, GIV’s GEM activity is induced upon capacitation. Once activated, GIV modulates both Gαi/s via the same GEM motif to suppress premature cAMP surges downstream of ligand-activated Gi/Gs-coupled GPCRs. Consequently, GIV-GEM inhibits premature acrosome shedding. Because both premature acrosome reaction or failure to do so are important causes of male infertility^55^, deciphering the signaling events that precisely regulate the timing of acrosome exocytosis has remained one of the most challenging and unresolved questions concerning mammalian reproductive biology^56^. Despite emerging evidence in the last decade that have challenged the long-held paradigms in the field, the identity of the pathways/processes/proteins that inhibit premature acrosome exocytosis was unknown. Our conclusion that GIV-GEM serves as a ‘brake’ for cAMP surge and prevents acrosome reaction is consistent with the fact that the PDE-inhibitor Sildenafil citrate (Viagra®) which increases cAMP causes premature acrosomal reaction^57^. It is noteworthy that although canonical G protein signaling that is triggered by ligand-activated GPCRs has been implicated in the activation/inhibition of mACs and cAMP signaling in the sperm head^58-60^, the role of non-canonical G protein we report here was never recognized previously. Because GIV is most highly expressed in sperm, the cAMP-regulatory role of GIV-GEM we define here implies that it may fulfil a major role in the regulation of cAMP in the sperm head. pYGIV was also detected in the sperm head, but its role in acrosome reaction was not studied here. Because pYGIV activates Class 1 PI3K, it is possible that the pYGIV→ PI3K axis at that location could also influence rapid lipid phosphorylations that are also known to regulate acrosome reaction^61^.

In the sperm tail, GIV is an effector within the sAC→ cAMP→ PKA→ multi-TK axis that gets robustly phosphorylated on Y1764; this site is known to directly bind and activate Class 1 PI3Ks, which ultimately enhance Akt signals. GIV’s GEM activity is also activated in the tails of capacitated sperms, and enhances Akt signals, presumably via the previously defined GIV→ Gi→ ’free’ Gβγ→ Class 1 PI3K axis. These two mechanisms of Akt signaling have previously been shown to act as “AND” gate to maximally enhance Akt signaling in diverse cell types to increase cell survival and motility^10,62^. Furthermore, both GIV transcripts and its phosphoactivation by TKs (pYGIV) was reduced in sperms with lower motility. Because the global trend of progressive reduction in the number of motile and viable sperm in the ejaculate has been associated with a concomitant increase in the rates of infertility^63^, our findings in the case of GIV add to the growing number of proteins that enrich the signal-ome of healthy sperm. For example, as an intrinsically disordered protein (IDP) and a multi-modular scaffold that generates crosstalk between diverse signaling pathways, GIV appears to be in a prominent position to orchestrate rapid cooperativity between these pathways and processes in the otherwise transcriptionally and translationally silent sperm cell.

In conclusion, our results provide evidence that GIV may perform different roles in the distinct spatial compartments of capacitating sperms. This study not only sheds light on defective GIV-signaling as potential ‘marker’ of male infertility, but also reveals that inhibitors of GIV-dependent signaling will inhibit fertility by reducing sperm motility and viability and by promoting premature acrosome reaction. The latter is a promising strategy for the development of a male contraceptive ‘pill’ specifically targeting sperm.

## ACKNOWLEDGEMENTS

We thank Masahide Takahashi, Ph.D (Nagoya University, Japan) for sharing Girdin fl/fl UBC-Cre-ERT2 mice and Lee Swanson for assisting with the initial breeding of the colonies. We thank members in the laboratory of Pamela L. Mellon, Ph.D. (UCSD) for helpful technical suggestions along the way. This work was supported by the National Institute of Health Grants: CA238042, CA100768, AI141630 and CA160911 (to Pr.Gh.), AI129894 and GM095882 (to Pa.Ga) and GM138385 (to DS). G.D.K was supported through The American Association of Immunologists Intersect Fellowship Program for Computational Scientists and Immunologists. I.L-S by the American Heart Association (AHA #14POST20050025) C.R was supported, in part, by an NIH-funded Training Grant Programs (T32 DK007202, T32 CA121938). This publication includes data generated at the UC San Diego IGM Genomics Center Utilizing an Illumina NOVASeq 6000 that was purchased with funding from a National Institutes of Health SIG grant (#S10 OD026929).

## AUTHOR CONTRIBUTIONS

S.R, in collaboration with I.L-S and C.R carried out and analyzed all the functional sperm assays in this work. V.C, C.R.E, and G.D.K performed all the mice fertility studies. S.T and D.S analyzed the transcriptomic datasets. Data visualization and statistical analyses were carried out by G.D.K. Pr.Gh. and Pa.Ga. P.G, conceptualized the studies, designed, supervised, and analyzed the experiments and wrote the first draft of the manuscript. G.D.K, Pr.Gh. and Pa.Ga. edited and revised the manuscript.

