## Supplementary Online Materials for "GIV/Girdin, a Non-receptor Modulator for Gαi/s, Regulates Spatiotemporal Signaling during Sperm Capacitation and is Required for Male Fertility"

**TRANSPARENT METHODS**

### Materials and methods

#### Human subjects

Human sperm were collected from healthy volunteers. A study proposal was approved by Institutional Review Board of University of California, San Diego. Isolation of human sperms were carried out as described under human research protocol (IRD #16027).

#### Animals

UbcreERT2+/- x Girdin fl/fl and Girdin fl/fl mice were generously provided by Dr. Takahashi (Nagoya University Graduate School of Medicine, Nagoya, Japan). Males UbcreERT2+/- x Girdin fl/fl were bred to females Girdin fl/fl to generate UbcreERT2+/- x Girdin fl/fl (experimental group), and UbcreERT2-/- x Girdin fl/fl (control group) mice. Genotyping was performed by PCR, and only male mice were used in this study. Wild type female C57/BL6 mice were purchased from The Jackson Laboratory (stock number: 000664; Bar Harbor, ME). All mice were housed in standard cages in an Association for Accreditation and Assessment of Laboratory Animal Care-approved animal facility at the University of California San Diego School of Medicine. This study was approved by the UCSD Institutional Animal Care and Use Committee, which serves to ensure that all federal guidelines concerning animal experimentation are met.

#### Reagents and antibodies

All reagents were of analytical grade and obtained from Sigma-Aldrich (St. Louis, MO) unless otherwise stated. The affinity purified anti-pS1689-GIV and pS1674-GIV were generated commercially in collaboration with 21st Century Biochemicals (Marlboro, MA) and validated previously[1,2]. Rabbit anti-GIV CT (T-13) was obtained from Santa Cruz Biotechnology; and a previously validated custom raised anti-p-GIV (pY1764) was from Spring Bioscience (Pleasanton, CA, USA)[3-6]. Mouse mAbs against pTyr was from BD Biosciences; Mouse anti-His, anti-a tubulin and anti-actin were obtained from Sigma; Rabbit monoclonal anti-Phospho-(p)Akt (Thr308) and anti-total (t)Akt were from Cell Signaling Technology (Beverly, MA). Rabbit anti-GIV-coiled-coil (CC) was obtained from EMD Millipore (Carlsbad, CA). Other commercially obtained antibodies used in this work were: Mouse anti-sp56 (MA1-10866) was purchased from Thermo Fisher Scientific (Waltham, MA), mouse Anti-Human Hexokinase 1/2 Monoclonal Antibody (Catalog # MAB8179; R&D Systems, Minneapolis, MN) and rabbit anti-phospho-PKA substrate (RRXS*/T*) (100G7E) mAb #9624 from Cell Signaling Technology.

Goat anti-rabbit and goat anti-mouse Alexa Fluor 680 or IRDye 800 F (ab0)2 used for immunoblotting were from Li-Cor Biosciences (Lincoln, NE). Goat anti-rabbit Alexa Fluor 488 and goat anti-mouse Alexa Fluor 594 for immunofluorescence were purchased from Life Technologies.

#### Immunohistochemistry of mouse testes

Mouse testes were fixed in zinc paraformaldehyde to prepare FFPE tissue blocks. Tissue sections of 4 µm thickness were cut and placed on glass slides coated with poly-L-lysine, followed by deparaffinization and hydration. Heat-induced epitope retrieval was performed using sodium citrate buffer (pH 6) in a pressure cooker. Tissue sections were incubated with 3% hydrogen peroxidase for 10 min to block endogenous peroxidase activity, followed by incubation with primary antibody overnight in a humidified chamber at 4°C. Antibodies used for immunostaining were SP173 rabbit monoclonal, anti-GIV antibody. Immunostaining was visualized with a labeled streptavidin–biotin using 3,3′-diaminobenzidine as a chromogen and counterstained with hematoxylin.

**Source of live sperms**

Mouse sperm suspension were obtained from cauda epididymis of mature male (9 weeks old) placed in 1 ml non capacitating (NC) buffer prewarmed at 38.1°C for 25 min. Sperm suspension allowed to incubate for 5 min and upper portion containing highly motile sperm were collected for the experiment.

Human ejaculates were collected from volunteers via masturbation after 1 week of abstinence under UCSD human subject protocol. After liquefaction at room temperature, 1 mL ejaculate was transferred into the bottom of 1 ml prewarmed NC buffer and incubated for additional 1 h to swim up procedure. Highly motile sperm mobilized to the upper layer was collected for the experiment.

***In vitro* capacitation and induction of the acrosome reaction**

Freshly obtained human and mouse sperms were segregated into low motile and high motile populations using ‘swim-up’ technique. Subsequently highly motile sperms were capacitated in TYH buffer containing 5 mg/ml BSA and 15mM $NaHCO_3$ at 37 °C under 5% CO2 for the indicated time mentioned in figure legends. Sperms were both lysed in reducing sample buffer for immunoblotting and fixed in 3% paraformaldehyde for immunofluorescence staining. Acrosomal reaction in sperm was triggered by incubating it either with Ca2+ ionophore or progesterone for the indicated time at 37 °C in 5% CO2. Sperm were then fixed and co-stained for peanut agglutinin (PNA-488; green, an acrosomal marker) and either pYGIV, or pSerGIV and DAPI.

**Confocal immunofluorescence**.

Sperms were fixed with 3% paraformaldehyde in PBS for 25 min at room temperature, treated with 0.1 M glycine for 10 min, and subsequently blocked/permeabilized with PBS containing 1% BSA and 0.1% Triton X-100 for 20 min at room temperature. Primary and secondary antibodies where incubated for 1 h at room temperature in PBS containing 1% BSA and 0.1% Triton X-100. Dilutions of antibodies used were as follows: GIV (1:500); phospho-GIV (Tyr1764; 1:500); phospho-pan-Tyr (1:500); α-tubulin (1:500); phosphor-GIV (Ser1674; 1:500); phosphor-GIV (Ser1689; 1:500); Peanut agglutinin (PNA) (1:500); His (1:500) and DAPI (1:2000). Secondary Alexa conjugated antibodies were used at 1:500 dilutions.

In the case of frozen sections of mouse testes, the protocol used was as follows: cryosections were washed three times with PBS, followed by 0.15% glycine for 10 min at room temperature and incubated for 20 min in blocking buffer (1% BSA in PBS), then 2 h in primary antibodies and 45 min in secondary antibodies. Dilutions of antibodies used were as follows: GIV (1:500); phospho-GIV (Tyr1764; 1:250); ZP3R (1:500); DAPI

(1:1000). Secondary Alexa conjugated antibodies were used at 1:250 dilutions. Sperms and sections were imaged on a Leica SPE confocal microscope using a 63x oil objective using 488, 561, 633 and 405 laser lines for excitation. The settings were optimized, and the final images scanned with line-averaging of 3. All images were processed using Image J software (NIH) and assembled for presentation using Photoshop and Illustrator software (Adobe).

**Dual color quantitative Immunoblotting**

Protein samples were separated by SDS/ PAGE and transferred to PVDF membranes (Millipore). Membranes were blocked with PBS supplemented with 5% nonfat milk (or with 5% BSA when probing for phosphorylated proteins) before incubation with primary antibodies. Infrared imaging with two-color detection and quantification were performed using a Li-Cor Odyssey imaging system. Primary antibodies were diluted as follows: anti-His 1:1,000; anti-GIV (tGIV) 1:500; anti–phospho-Tyr-1764-GIV (pYGIV) 1:500; anti-phospho-Tyr (pan pY) 1: 500; anti-phospho-PKA 1: 500; anti-Hexokinase 1:1000; anti-phospho-Akt (Thr308) 1:500; anti-Akt 1:500; anti-β tubulin 1:1000. All Odyssey images were processed using Image J software (NIH) and assembled for presentation using Photoshop and Illustrator software (Adobe).

**His-TAT purification and transduction in sperms.**

Cloning of TAT-GIV-CT-WT and TAT-GIV-CT-FA mutant has been described[7]. TAT-constructs were expressed using BL21(DE3)-pLysS (Invitrogen) and Terrific Broth (BioPioneer) supplemented with additives as per auto-induction protocols outlined by Studier F[8]. Briefly, cultures of bacteria were grown at 300 rpm at 37°C for 5 h, then at 25°C overnight. Cells were lysed in 10 mL of lysis buffer containing 20 mM Tris, 10 mM Imidazole, 400 mM NaCl, 1% (vol:vol) Sarkosyl, 1% (vol:vol) Triton X-100, 2 mM DTT, 2 mM Na3oV4 and protease inhibitor mixture (Roche Diagnostics), pH 7.4, sonicated (3 × 30 s), cleared at 12,000 × g for 20 min at 4°C and affinity-purified on Ni-NTA agarose resin (Qiagen) (4 h at 4 °C). Proteins were eluted in elution buffer containing 20 mM Tris, 300 mM Imidazole, 400 mM NaCl, pH 7.4, dialyzed overnight against TBS containing 400 mM NaCl and stored at −80 °C.

TAT transduction in sperms was performed by incubating them with 400–800 nM TAT-GIV-CT peptides for 30 min. Efficiency uptake was measured by flow cytometry.

**CASA system**

Freshly obtained sperms were segregated into low motile and high motile populations using ‘swim-up’ technique and highly motile sperms were subsequently capacitated in TYH buffer containing 5 mg/ml BSA and 15mM $NaHCO_3$ along with TAT-GIV-CT peptides at 37 °C under 5% CO2 for 3h. The sperm motility and progressive motility were measured on CASA on a Hamilton Thorne IVSO-CASA (Berns Laboratory, UC San Diego).

**Measurement of sperm cAMP level**

Mouse sperms at a density of 2 x $10^7$ cells/ml (6 x $10^6$ cells in total) were first peptide transduced with TAT-GIV-CT for 30 min, washed gently with PBS three times to remove excess peptides before their use in cAMP assays.

Peptide-transduced sperms were pre-incubated with 0.5 mM isobutyl methyl xanthine (IBMX) prior to exposure to various chemicals at the following final concentrations: 25 mM $NaHCO_3$, 0.1 mM adenosine, or 100 μM After mixing with the respective stimulus, the samples were incubated for 30 min at 37°C, followed by the addition of 0.25 M HCl (final concentration) to quench the biochemical reactions. After incubation for 30 min at room temperature, cell debris was sedimented by centrifugation at 3,000 g for 5 min at room temperature. The cAMP concentration in the supernatant was determined by a competitive enzyme immunoassay according to the product manual (Catalog #: ADI-900-066, Enzo Life Sciences).

**Tamoxifen treatment and Natural mating**

4–5-week-old mice experimental or control mice received an intraperitoneal (i.p.) tamoxifen injection 1 mg/100ul/mouse/day (Millipore Sigma, St. Louis, MO) solubilized in 100% corn oil for five consecutive days. Girdin gene knockout (GIV-knockout) was confirmed using DNA qPCR as described in our previous study [9]. Mice were then housed for 3 weeks with random females to promote mating with the intent to discharge sperms in which tamoxifen has not yet induced Cre expression.

After 3 weeks, each male mouse was housed with three 7-8 weeks old C57BL/6 fertile females for 3 months. The frequency of successful live births and the litter size were recorded. After 3 months, experimental and control males were sacrificed, and testicles and epididymis were collected for Immunohistochemistry (IHC), immunoblotting and mRNA analysis.

**Transcriptomic datasets from infertile patients**

Publicly available RNASeq dataset (GSE4797[10], E-TABM-234[11], GSE6872[12], GSE103905[13] and GSE26881[14]) were downloaded from National Center for Biotechnology Information (NCBI) Gene Expression Omnibus website (GEO) and ArrayExpress. The data was processed using the Hegemon data analysis framework repositories for male infertility[15-17]. All semen samples mentioned in above datasets were classified based WHO (2010) guidelines for semen parameters[18].

| WHO (2010) guidelines[18] (Semen parameters) | Motility (%) | Morphology (%) | Concentration ($10^6$/mL) |
|---|---|---|---|
| Fertile individual | $\geq 40$ | $\geq 4$ | $\geq 15$ |

**Total RNA isolation**

Total RNA from mouse testicles were isolated using the Direct-zol RNA Miniprep Kit (Zymo Research, Irvine, CA) following manufacturer's instructions. RNA concentration was measured using the NanoDrop™ One$^C$ (Thermo Scientific, Waltham, Massachusetts). The RNA Integrity Number (RIN) was assessed using the 4200 TapeStation system and the TapeStation RNA ScreenTape & Reagents (Agilent Technologies, Santa Clara, CA).

**RNA-seq and data analysis**

Total RNA samples were submitted to the IGM Genomics Center (University of California San Diego) for library preparation and sequencing. mRNA stranded sequencing libraries were generated with the TruSeq Stranded mRNA Sample Prep Kit with TruSeq Unique Dual Indexes (Illumina, San Diego, CA). Resulting libraries were multiplexed and sequenced with 100-bp paired-end reads (PE100) to a depth of approximately 30 million reads per sample on an Illumina NovaSeq 6000. Samples were demultiplexed using bcl2fastq v2.20 Conversion Software (Illumina, San Diego, CA).

To determine, which genes were differentially expressed in WT and GIV-knockout mice testes, transcript-level abundance of paired-end RNA-seq data was estimated by Salmon (1.1.0) using the mouse transcriptome from Genecode (vM24). Tximport (1.14.2) is used to aggregate transcript-level quantification to the gene-level. The resulting gene counts were used as an input to DESeq2 Bioconductor package. Differentially expressed genes below a BH adjusted p-value of 0.05 were considered significant. Also, all genes differentially expressed were included in REACTOME pathway enrichment analysis. Statistically significant pathways of up-regulated and down-regulated DEGs are listed in the table and bar plots of up-regulated and down-regulated enriched pathways. In GSEA analysis, some fertility-related gene sets from Molecular Signatures Database (MSigDB) were tested. Genes from these gene sets were used to rank order the samples and test for GIV-knockout versus WT phenotype classification using the AUC (Area Under the Curve) ROC (Receiver Operating Characteristics) curve and displayed such classification using violin plots.

**Statistical Analysis**

Statistical significance between datasets with three or more experimental groups was determined using one-way analysis of variance (ANOVA) followed by Tukey’s test for multiple comparisons. Unpaired t-test is used to test the statistical difference between two experimental groups. For all tests, a p-value > 0.05 is considered as significant. All experiments were repeated a least three times. All statistical analysis was performed using GraphPad prism 9.

**Data and Software Availability**

RNA Seq is uploaded to NCBI GEO [GSE171704].

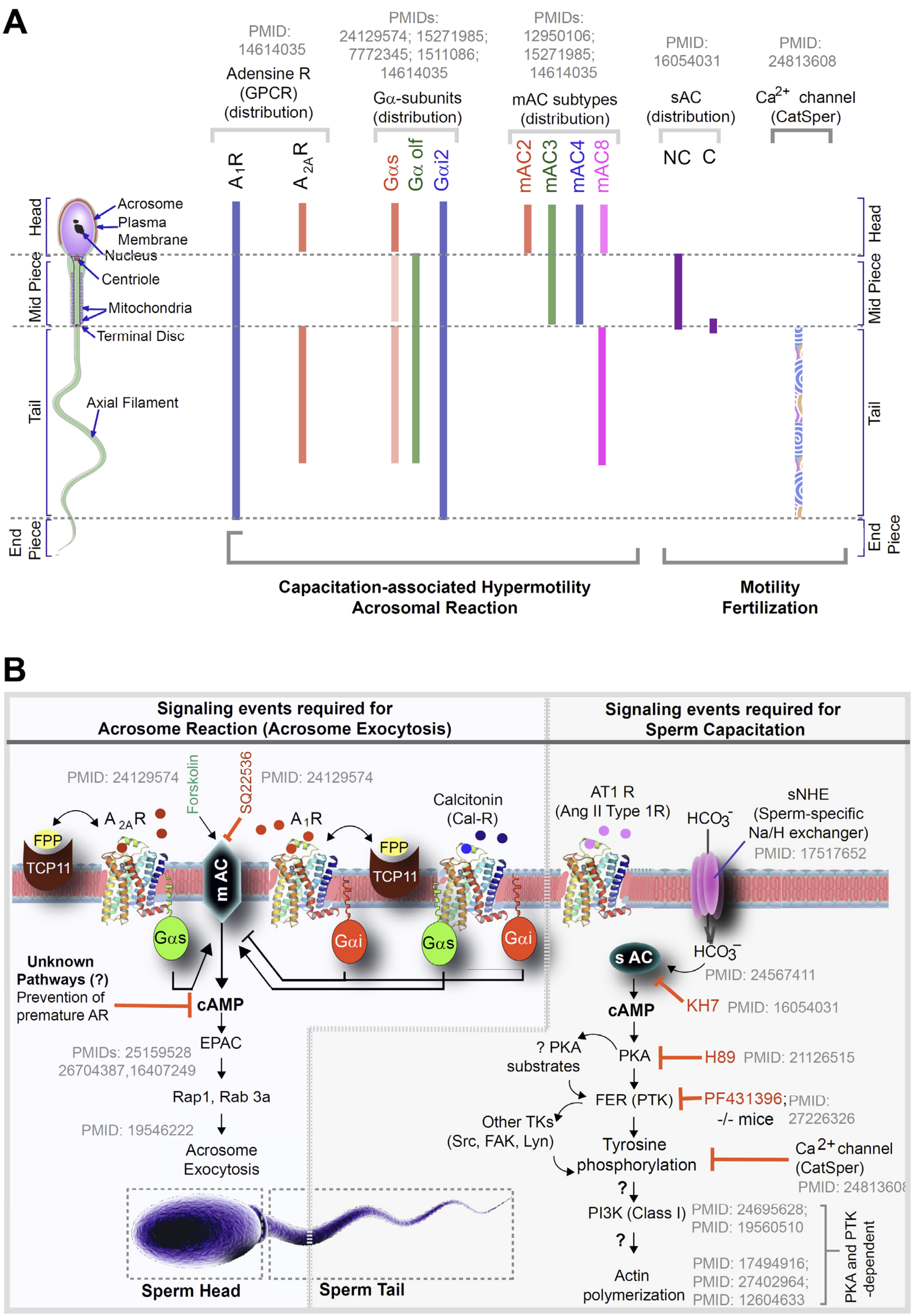


**Figure 1- Figure Supplement 1**
**Schematic summarizing the known localization (A) and role of G proteins/AC proteins and their impact on cAMP signaling (B) during sperm processes.** A. The localization of G protein subunits and membrane and soluble adenylyl cyclases (mAC/sAC) and the calcium channel, Catsper is shown. The intensity of the bar denotes the relative concentrations of the molecules. B. Summary of experimental evidence and key citations and unknown (?) aspects in spatiotemporally separated signaling cascades that regulate sperm processes.

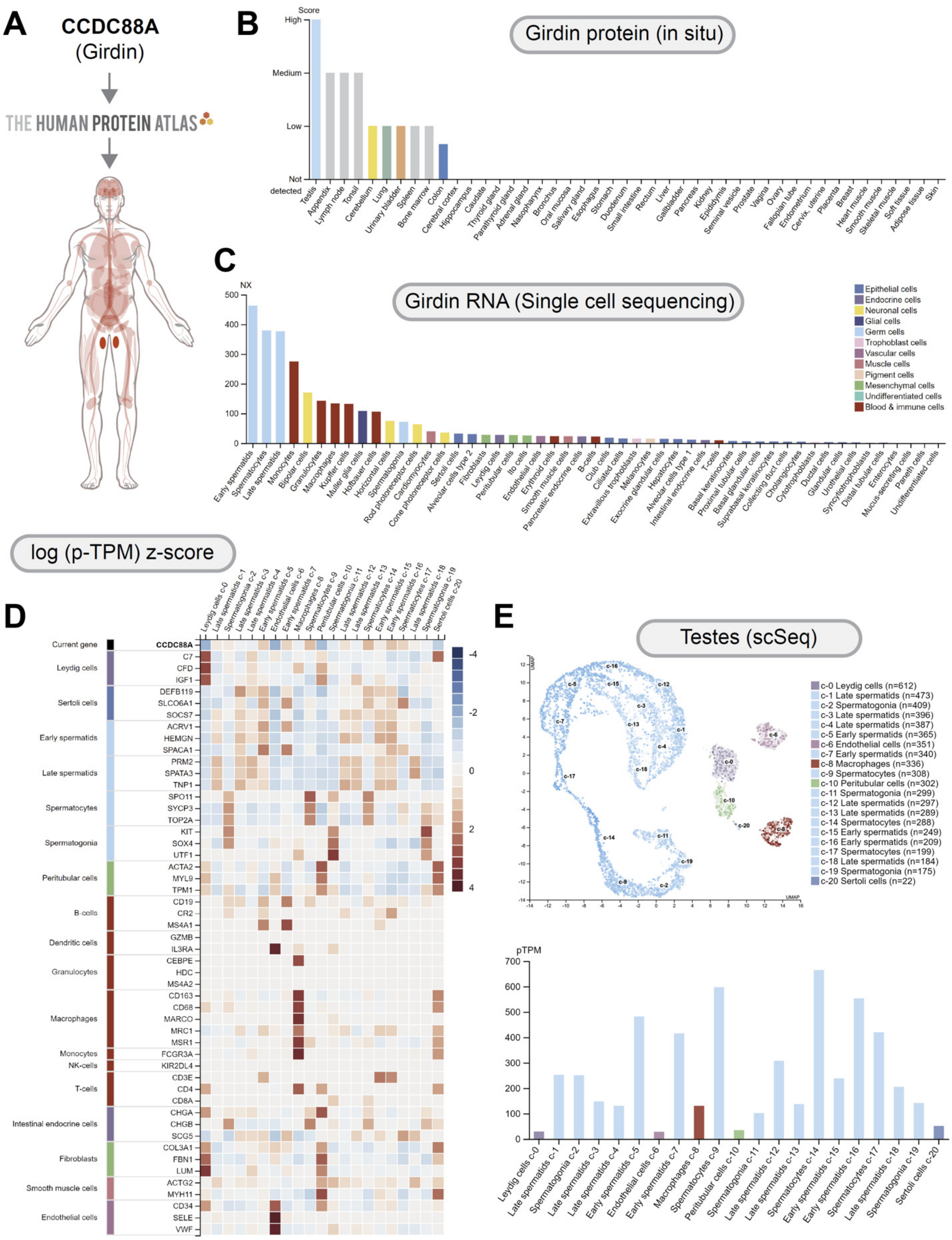


**Figure 1- Figure Supplement 2**

**CCCD88A (GIV/Girdin) is highly expressed in the testes, most specifically the spermatocytes.**

**A**. CCDC88A expression profile was queried in the Human Protein Atlas (HPA). **B.** GIV protein expression data is shown for each of the 44 tissues. **C**. A summary of single cell RNA (NX) from all single cell types. Color-coding is based on cell type groups, each consisting of cell types with functional features in common. **D**. The heatmap in this section shows expression of CCDC88A and well-known cell type markers in the different single cell type clusters of human testes. Normalized data is presented as log (p-TPM) Z-score. **E**. UMAP PLOT (top) for single cell expression of CCDC88A in human testes. A bar chart (bottom) shows RNA expression (pTPM) in each cell type cluster.

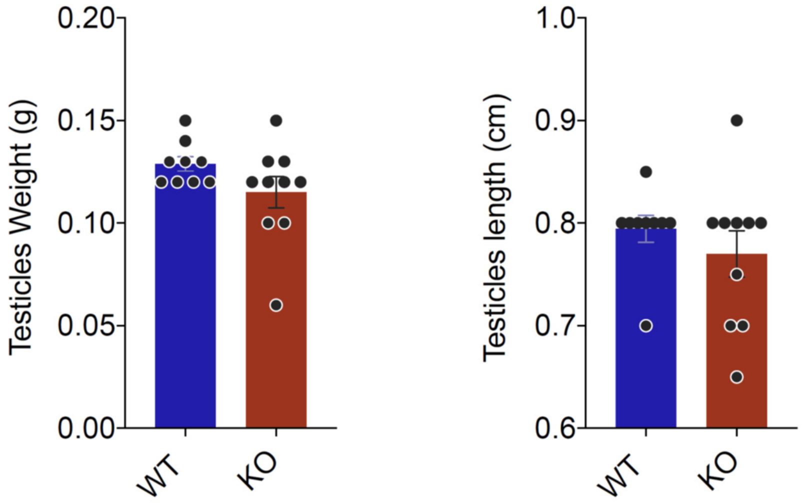


**Figure 5- Figure Supplement 1**
**GIV is required for male fertility but its depletion does not impact testes weight or length.**
GIV was depleted in male mice by intraperitoneal injection of Tamoxifen (see Figure 5A). Bar graphs show the testes weight (Left) and length (Right) in WT and GIV-KO males.

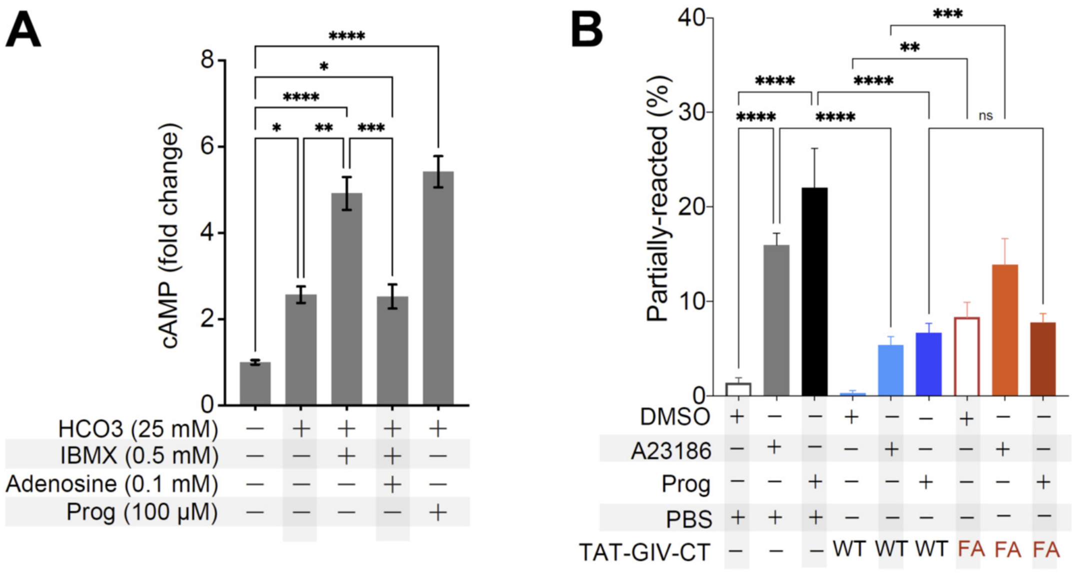


**Figure 7- Figure Supplement 1**
**An intact GEM motif in GIV is required for inhibiting cAMP surge and acrosomal reaction**
**A.** Bar graphs display the fold change in cAMP in mouse sperms in the presence (+) or absence (-) of various treatments. **B**. Acrosomal reaction was analyzed in TAT-GIVCT transduced mouse sperms exposed to either DMSO control or the calcium ionophore A23186 or Progesterone using CD46 as a marker of inner acrosomal membrane (IAM) as outlined in Figure 7D. Bar graph presented here shows the proportion of partially reacted sperms in each treatment group.
